# Epigenetic lockdown of type I interferon sensing and signalling in human pluripotent cells

**DOI:** 10.1101/2025.10.27.684762

**Authors:** James H. Holt, Rocio Enriquez-Gasca, Rachel P. Wilson, Elena G. Bochukova, Pierre V. Maillard, Helen M. Rowe

**Affiliations:** Centre for Immunobiology and Infection, Blizard Institute, Queen Mary University of London, E1 2AT, UK

**Author notes:** The Francis Crick Institute, 1 Midland Road, London NW1 1AT, UK. Contributed equally (Pierre V. Maillard, Helen M. Rowe).

## Abstract

The Human Silencing Hub (HUSH) complex safeguards genome integrity in human somatic cells by repressing transposable elements and regulating type I interferon (IFN-I) induction. In early development, the IFN-I pathway is inactive, yet its underlying regulation is poorly understood. Here, we use depletion of the HUSH complex in human induced pluripotent stem cells (iPSCs) as a tool to investigate epigenetic control of the IFN-I system in early development. We confirmed that human iPSCs display an attenuated IFN-I pathway, whereas iPSC-derived neural progenitor cells (NPCs) respond robustly to IFN-I pathway agonists. We found that, in iPSCs, depletion of MPP8, a core component of all HUSH complexes, was sufficient to induce both expression of young LINE-1 elements and genes linked to the IFN system including double-stranded RNA sensors and interferon-stimulated genes (ISGs). ISG upregulation had little effect on pluripotency markers and occurred without IFN signalling, suggesting that, in contrast to differentiated cells, these ISGs are direct transcriptional targets of the HUSH complex in early development. Chromatin profiling by CUT&Tag confirmed MPP8 enrichment at HUSH-regulated ISGs and revealed a bimodal binding profile of MPP8 to both ISGs and non-ISGs, the latter largely driven by young LINE-1 elements. We propose that shutdown of the IFN-I system in pluripotent stem cells is essential to prevent lethality from unwarranted self-nucleic acid sensing. This shutdown is achieved through a triple-layer of epigenetic lockdown acting on ligands, sensors, and effectors across the IFN-I pathway. Pluripotent cells, therefore, represent a ground state of immune evasion that cancer cells may evolve towards through increasing expression of MPP8.

## Introduction

A large proportion of the noncoding genome is made up of transposable elements (TEs), which are parasitic sequences that can change their location or increase their copy number within the genome ^1–3^. Although most TEs have lost mobility due to accumulated mutations, many retain promoter or enhancer activity and can still influence host gene regulation. One group of TEs of particular interest are Long INterspersed Element-1s (LINE-1s) ^4–8^. The youngest LINE-1 subfamily in humans, L1HS (also known as L1PA1), is unique to hominids and represents the only remaining mobile and autonomous retrotransposon subfamily in humans. L1HS elements can also mediate the transposition of non-autonomous retrotransposons, including older LINE-1s, short interspersed nuclear elements (SINEs) and SINE-VNTR-Alu (SVA) elements in *trans* ^9^. The human genome contains approximately 8,000 full-length LINE-1 copies, along with numerous truncated fragments, together accounting for around 17% of the genome ^2^. The links between TE activity and disease are well established with particularly strong associations to genetic diseases ^10–12^ and cancer ^13–15^. Yet, beyond these deleterious effects, TEs are also frequently co-opted to serve regulatory roles in our genome. They can modulate gene expression without retrotransposition through diverse mechanisms including controlling cell type-specific gene transcription ^16^, acting as enhancers ^17–19^, generating novel isotypes via exonisation of genes ^20^, contributing to long noncoding RNA (lncRNA) ^21^, and even giving rise to entirely novel genes ^22,23^.

Given the risks associated with TEs, their activity is tightly restrained by multiple mechanisms of epigenetic repression including DNA methylation ^24–26^, histone modifications ^27,28^, changes in chromatin structure ^29^, and post-transcriptional silencing ^8^. These, in turn, are controlled by an array of epigenetic complexes including Krüppel-associated box domain zinc finger proteins (KRAB-ZFPs) and the Human Silencing Hub (HUSH) complex ^30–32^. The core HUSH complex (also called HUSH1) consists of MPP8, Periphilin (PPHLN1) and TASOR alongside an array of accessory proteins. Its recruitment relies on several mechanisms. The chromodomain of MPP8 binds H3K9me3 ^32^, while PPHLN1 associates with specific RNA transcripts including RNAs from young LINE-1s, processed pseudogenes and adenosine-rich intronless genes ^33^. ZNF638 (NP220) interacts specifically with TASOR and has been implicated in recruiting HUSH to transposable elements and non-integrated viral DNA ^34–36^. In addition, the pseudo-PARP domain of TASOR has been shown to play a critical role in the localisation of HUSH to LINE-1 elements ^37^. HUSH-mediated silencing also depends on the recruitment of various effector proteins and complexes including MORC2 ^7,31^, SETDB1, and ATF7IP ^38,39^. In addition, HUSH targets TE-derived RNAs through interactions with the nuclear exosome targeting (NEXT) complex ^40,41^, and with the poly-(A) degrading protein CNOT1 ^41^.

In mammals, the type I interferon (IFN-I) system is a central defence pathway against viral invasion ^42–44^. It is initiated when innate immune receptors recognise viral nucleic acids such as double-stranded RNA (dsRNA), which often accumulates during viral replication. Receptor activation initiates a signalling cascade that activates interferon regulatory factors (IRFs) to drive transcription of IFNs, primarily IFNα and IFNβ. These cytokines are secreted and bind the IFNα/β receptor (IFNAR) in a paracrine or autocrine manner, activating JAK/STAT signalling. This leads to the formation of the interferon stimulated gene factor 3 (ISGF3) complex, which binds IFN-stimulated response elements (ISREs) in target gene promoters to induce expression of hundreds of interferon stimulated genes (ISGs). While essential for antiviral protection, aberrant activation by self-derived nucleic acids has been reported in autoinflammatory and autoimmune diseases as well as in cancer ^45^. It is therefore crucial for this response to be tightly regulated and activated only upon infection.

While the IFN-I response constitutes a key defence mechanism against viral infection in most cells, it remains inactive in pluripotent stem cells and during early development. Early studies showed that mouse embryonal carcinoma (EC) cells and mouse embryonic stem cells (mESCs) were refractory to IFN induction following viral infections and exposure to dsRNA ^46–48^. Likewise, early embryos are incapable of producing IFN and their ability to synthesise IFN occurs from embryonic day 7 ^46–48^. A similarly inactive IFN induction was also observed in human ESCs as well in human induced pluripotent stem cells (iPSCs), which are both also unresponsive to IFN treatment ^49–52^. Interestingly, *in vitro* differentiation of mESCs is accompanied by a gain of IFN expression upon virus exposure ^53^. Conversely, reprogramming of fibroblasts, which possess a robust IFN-I response, into hiPSCs results in a loss of IFN expression after viral challenge, suggesting a reciprocal inhibition between pluripotency and the IFN system ^54^. Unable to either produce or respond to IFNs, stem cells instead rely on alternate mechanisms such as RNA interference and the intrinsic expression of a subset of ISGs to protect against viruses ^50,55–58^. It has been proposed that these differences in antiviral defence by stem cells may be due to their unique requirements. Many ISGs have well-documented antiproliferative and pro-differentiation effects, acceptable in differentiated cells but detrimental to stem cells and early development ^59^. Consistent with this notion, artificial induction of the IFN-I system in iPSCs was seen to impair their pluripotency and ability to differentiate ^51^. We therefore hypothesise that an uncharacterised mechanism operates in stem cells to lock down the IFN-I system.

We previously demonstrated that loss of MPP8 in differentiated primary fibroblasts causes expression of LINE-1 RNA, including overlapping sense/antisense RNAs capable of forming dsRNA ^60^. MPP8 depletion led to the accumulation of endogenous dsRNA and induced the expression of IFNβ and ISGs in an IFNAR-dependent manner and dependent on sensing via the dsRNA sensors RIG-I and MDA5 ^60^. Recent work has identified an alternate version of the HUSH complex, named HUSH2, which shares MPP8 and PPHLN1 but incorporates TASOR2, a paralogue of TASOR. Unlike HUSH, which primarily represses transposable elements ^61,62^, HUSH2 regulates the expression of certain ISGs ^63,64^. HUSH2 localizes to specific gene promoters and enhancers including those of ISGs and KRAB-ZNF genes with recruitment to ISGs mediated by TASOR2, which binds at IRF1, IRF2, and ISRE motifs ^36,65,66^. The mechanisms of HUSH2-mediated silencing are less well characterised. TASOR binding sites overlap with repressive H3K9me3 modifications and not active H3K4me3 whereas TASOR2 sites frequently coincide with H3K4me3 modifications and not H3K9me3 ^67^, suggesting that HUSH2 silences genes via a mechanism distinct from HUSH. Interestingly, it was found that TASOR and TASOR2 exist in balance where TASOR depletion or increased expression of TASOR2 increases TE expression, while TASOR2 depletion elevates ISG expression ^36^. In sum, the HUSH complex regulates a complex web of epigenetic and RNA-targeting mechanisms to silence both TEs and genes.

In this work, we employed the HUSH complex to investigate if and how epigenetic silencing regulates the IFN-I system in early human development, which represents an immune privileged stage. Our results reveal that epigenetic lockdown of the IFN-I system ensues in early development and acts at three levels to switch off ligands, sensors, and effectors of the IFN-I system.

## Results

### iPSCs display significantly lower sensitivity to IFNβ and dsRNA agonists than isogenic NPCs

To confirm previous reports that iPSCs exhibit a relative inability to mount an IFN response, we tested a panel of agonists: two dsRNA sensing agonists of RIG-I and MDA5 (the dsRNA analogue poly(I:C) and a 5’-triphosphate hairpin RNA (3p-hpRNA)) as well as to recombinant IFNβ, in iPSCs (iPSC 2), and iPSCs differentiated into NPCs, the latter having been shown to mount effective IFN responses ^55^ by monitoring changes in expression of canonical ISGs (i.e. IFIT1, IFIT2 and MX1) (Figure 1A, Supplementary Figure 1A). In iPSCs, ISG induction was relatively modest (from 1.5 to 13-fold), with IFIT2 showing the highest increase following poly(I:C) treatment (Figure 1B). In contrast, NPCs exhibited a markedly stronger response with all three ISGs upregulated from ∼20 to ∼3300 fold and MX1 showing the strongest induction after poly(I:C) treatment (Figure 1B). Therefore, we found that upregulation of ISGs was much higher in NPCs than iPSCs following treatment with IFNβ, poly(I:C), or 3p-hpRNA, indicating that our iPSCs have an attenuated IFN response compared to their differentiated NPCs counterparts.

**Figure 1:**
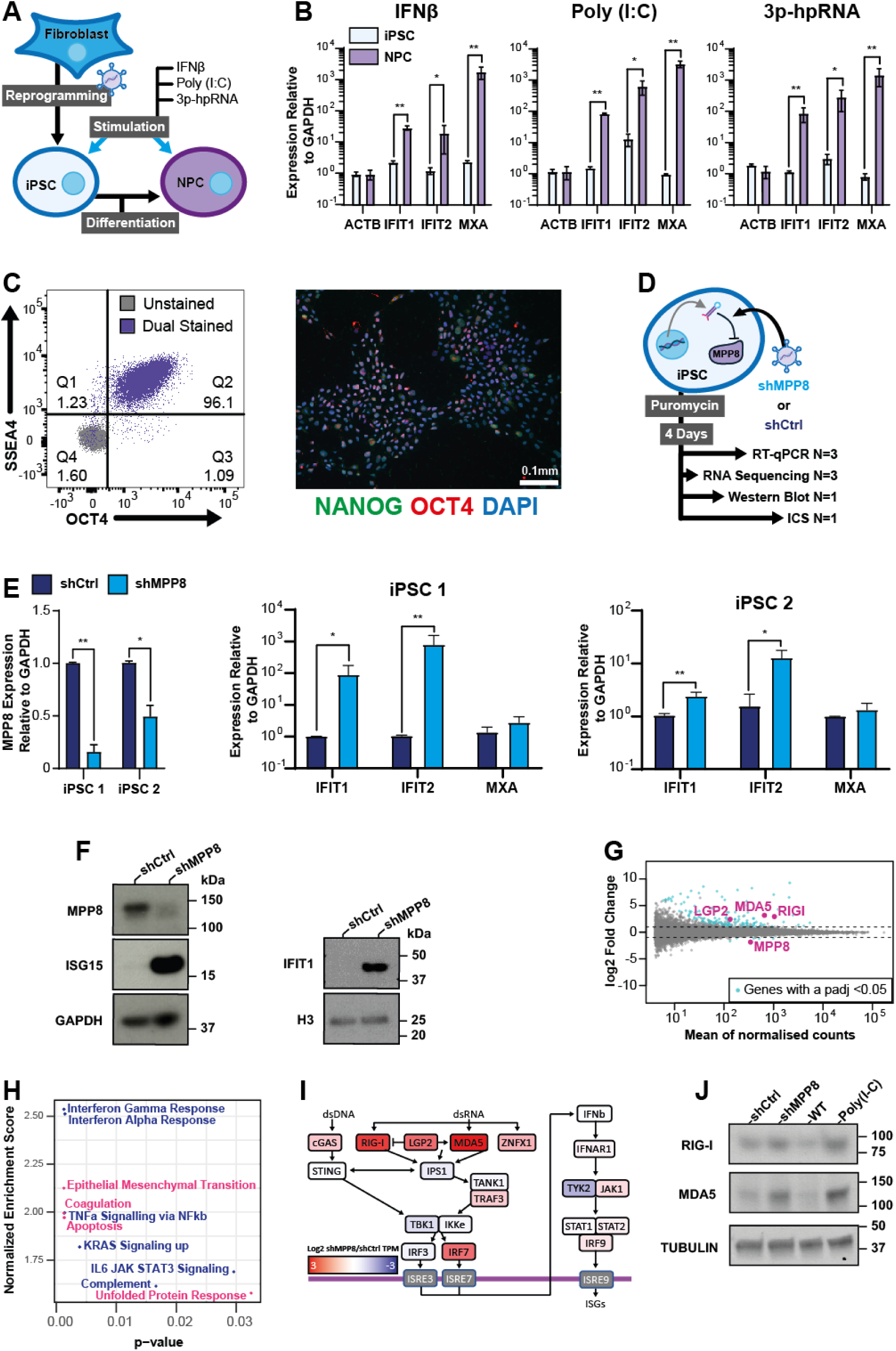
MPP8 depletion in iPSCs induces ISG expression and restores the expression of key nucleic acid sensors. (A) Schematic showing iPSC reprogramming followed by differentiation into NPCs and subsequent treatment with agonists of dsRNA sensing and IFN signalling. (B) RT-qPCR showing ISG expression relative to GAPDH in iPSCs (blue) and NPCs (purple) after treatment with IFNβ (left), poly(I:C) (middle), and 3p-hpRNA (right). (C) validation of iPSC expression of pluripotency markers by intracellular staining (ICS) followed by flow cytometry (iPSC 1, left) or microscopy (iPSC 2, right). (D) Schematic of iPSC transduction with lentiviral vectors expressing either a non-targeting shRNA control (shCtrl) or a MPP8-specific targeting shRNA (shMPP8) showing the number of replicates analysed using each technique. (E) RT-qPCR showing MPP8 depletion (left) or ISG upregulation (middle, right) relative to GAPDH in iPSC 1 and iPSC 2 after shRNA treatment. (F) Western blots on extracts from iPSC 1 following MPP8 depletion probed for MPP8, ISG15, and IFIT1 with GAPDH and Histone H3 serving as a loading control. (G) Annotated MA plot showing results of differential expression analysis following MPP8 depletion. 149 significantly differentially expressed genes (log2 FC > 1 and a p-adjusted value < 0.05 after Benjamini– Hochberg multiple testing correction of Wald test p-value of shMPP8/shCtrl) are shown in light blue while MPP8 and genes related to dsRNA sensing highlighted in pink. (H) Enrichment plot of significantly enriched gene sets from three biological replicates, colour coded for those common to previously shown in human foreskin fibroblasts (HFFs) (blue) and those unique to iPSCs (pink). (I) Colour-coded pathway diagram indicating log2 FC of genes involved in nucleic acid sensing following MPP8 depletion in iPSCs. (J) Western blots on extracts from iPSC 1 following MPP8 depletion probed for RIG-I and MDA5 with GAPDH as a loading control. Bar graphs show N = 3 biologically independent samples with SD shown as error bars (B, E), and BH-adjusted p-values from two-tailed paired t tests (B) or unadjusted p values from two-tailed unpaired t tests (E). ** = P < 0.01 and * = P < 0.05.

### MPP8 depletion in iPSCs induces ISG expression and restores the expression of key nucleic acid sensors

To investigate how the HUSH complex regulates ISGs in stem cells we used two different iPSC lines: WIBJ2 (iPSC 1) and WTC11 (iPSC 2). To validate that these cells express pluripotency markers, we used intracellular flow cytometry to stain for OCT4 and SSEA4 in iPSC 1 (Figure 1D, 1C, left, Supplementary Figure 1B). In iPSC 2, we used microscopy to validate OCT4 and NANOG expression (Figure 1C, right, Supplementary Figures 2A-G), as well as RT-qPCR for NANOG and OCT4 (Supplementary Figure 1C), and Western blot for OCT4 (Supplementary Figure 1D). Having validated that both cell lines were positive for pluripotency markers, we depleted MPP8 using short hairpin RNAs (shRNAs) compared to a non-targeting control (shCtrl) and selected the transduced cells with puromycin for four days before analysis (Figure 1C).

We documented effective depletion of MPP8 in both iPSC 1 (84%) and iPSC 2 (50%) (Figure 1E, left). In both iPSC lines we then measured expression of a panel of ISGs previously shown to respond to MPP8 depletion in human foreskin fibroblasts (HFFs) ^68^. Expression of IFIT1 and IFIT2 was markedly increased in both cell lines with a much stronger upregulation in iPSC 1 (88-fold and 781-fold, respectively) compared to iPSC 2 (2.4-fold and 13-fold) (Figure 1E). iPSC 1 exhibited both stronger MPP8 depletion and ISG upregulation suggesting a dose-dependent response. As MPP8 depletion was more effective in iPSC 1, subsequent experiments were carried out in this line. Western blotting confirmed that OCT4 expression was restricted to iPSC 1 compared to somatic cell lines (Supplementary Figure 1D) and demonstrated that MPP8 depletion increased protein levels of the ISGs, IFIT1 and ISG15 (Figure 1F).

To broadly assess the impact of MPP8 depletion on genes and TEs in iPSCs, we performed total RNA sequencing (RNA-seq). Differential expression analysis revealed downregulation of six genes, including MPP8, and upregulation of 143 genes (Figure 1G). Notably, seven of the top ten most strongly upregulated genes were ISGs.

We then used gene set enrichment analyses to identify pathways affected by MPP8 depletion in iPSCs. Many enriched gene sets were common with those previously reported in HFFs following MPP8 depletion, including IFNα targets, IFNγ targets, TNFα, KRAS signalling, IL6-JAK/STAT signalling, and activation of complement pathways (Figure 1H) ^68^. Intriguingly, upregulated gene sets unique to iPSCs included epithelial-mesenchymal transition (EMT), coagulation, apoptosis, and the unfolded protein response. Full lists of genes and gene sets are available in the source data.

Further investigation of the dsRNA sensing pathway showed that several key components of this pathway, including RIG-I and MDA5, are silenced by MPP8 at both the mRNA (Figure 1I) and protein levels (Figure 1J). Many components of the dsRNA sensing pathway were found to be expressed only at low levels in shCtrl iPSCs (Supplementary Figure 1F), consistent with the inactivity of the IFN-I system in iPSCs.

### MPP8 depletion in iPSCs induces expression of young, full-length LINE-1 elements

To investigate the effects of MPP8 depletion on TE expression in iPSCs at the subfamily level we used Tecount ^69^ with multimapping reads and DESEQ2 ^70^ to identify differentially expressed TE subfamilies. We found that six TE subfamilies were significantly upregulated following MPP8 depletion (Figure 2A). These included the youngest three subfamilies of LINE-1s (L1HS, L1PA2, L1PA3) as well as 2 LTR families (MER9A and LTR1D) and a DNA transposon family (HSMAR1). Comparison of the relative ages of these TEs compared to primate evolution revealed that these subfamilies were all relatively young, with the oldest, LTR1D, being shared by all Haplorrhini primates, which diverged from non-Haplorrhini primates approximately 70 million years ago (Figure 2B). The oldest of the LINE-1 subfamilies identified, L1PA3 is common to all Homininae (a Subfamily including humans, gorillas, and chimpanzees) and has been estimated to be approximately 12.5 million years old ^71^.

**Figure 2:**
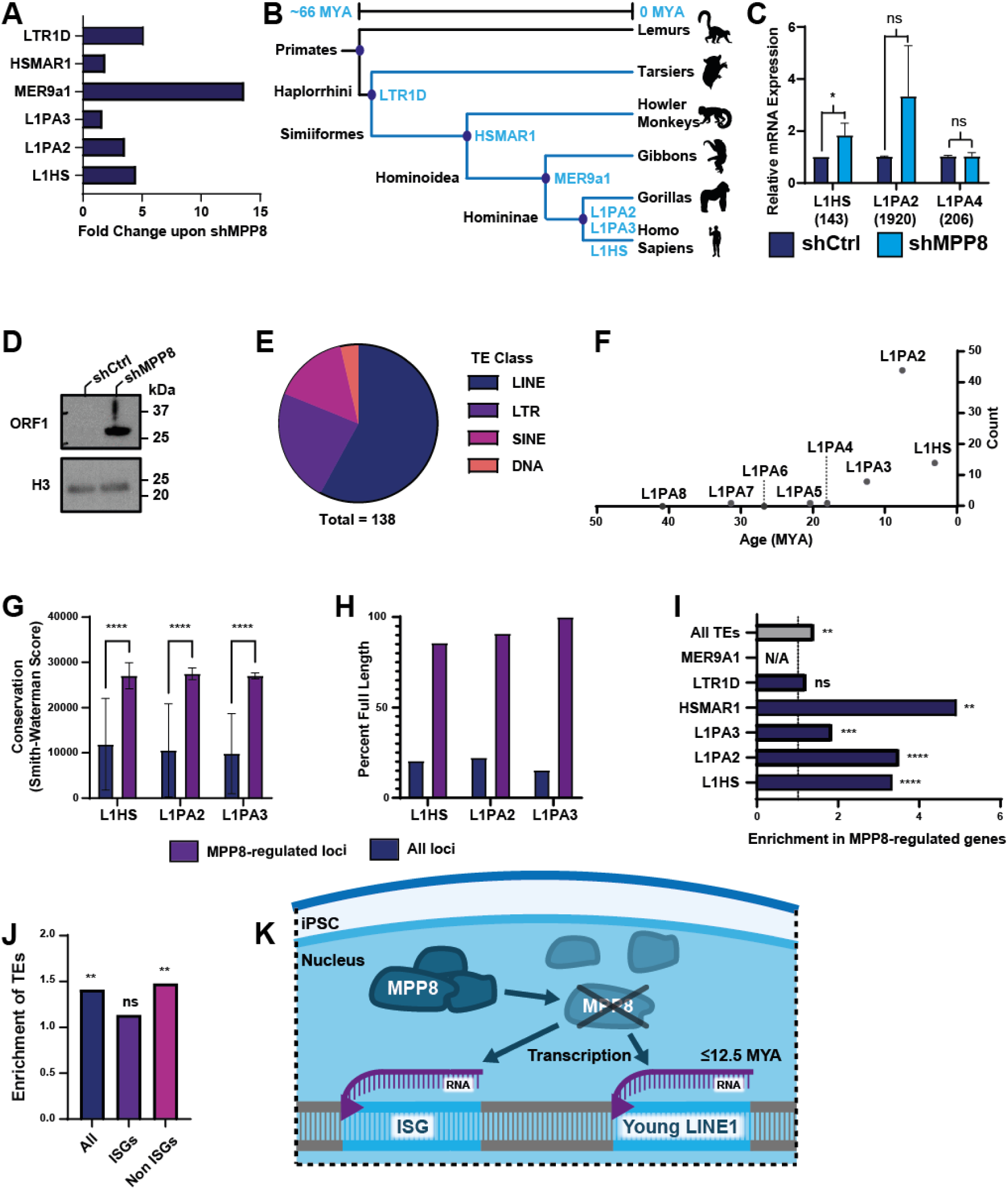
MPP8 depletion in iPSCs induces expression of young, full-length LINE-1 elements. Total RNA sequencing was used to identify variation in TE expression with three independent biological replicates. (A) Fold change of differentially expressed TE subfamilies as identified by Tecount. (B) Representative phylogenetic tree of the Order primates showing modern examples of each Suborder, Infraorder, Family, and Subfamily annotated with TE subfamilies (light blue) from (A) where they first appear as per DFAM. (C) RT-qPCR quantifying TE expression using primers targeting 143 L1HS loci, 1920 L1PA2 loci, and 206 L1PA4 loci, all shown relative to GAPDH in shCtrl-expressing cells. (D) Western blot on extracts from iPSC 1 expressing shCtrl or shMPP8 probed for LINE-1 ORF1 and Histone H3 as a loading control. (E) Pie chart showing the TE classes of 138 differentially expressed TE loci identified by SQuIRE following MPP8 depletion. LINEs are shown in blue, LTRs in purple, SINEs in pink and DNA transposons in orange. (F) Scatter plot showing the age of LINE-1 subfamilies as per Khan et al, 2006 and the number of upregulated loci of that subfamily identified by SQuIRE. (G) Average Smith-Waterman Score (SWS) of upregulated loci (purple) versus all loci of that subfamily (blue) for L1HS, L1PA2, and L1PA3, annotated with p-values from two-tailed unpaired t tests. **** = P < 0.0001 (H) Bar plot showing the percentage of upregulated LINE-1 loci that are full-length (Repeatmasker annotation length between 5.5kb and 6.5kb, purple) versus all loci of that subfamily (blue) for L1HS, L1PA2, and L1PA3. (I) Bar plot showing enrichment of all TEs (grey) and MPP8-regulated TE subfamilies (blue) in genes differently expressed upon MPP8 depletion. Enrichment was calculated for the 143 MPP8-regulated genes compared with 10,000 shuffled groups of 143 genes. No overlaps between MPP8-regulated genes and MER9A1 were found so no enrichment could be calculated. (J) Bar plot showing enrichment of all TEs in all 143 MPP8-regulated genes (blue) compared with MPP8-regulated ISGs and non-ISGs. P-values were calculated based on the number of shuffled gene sets with more overlap than our MPP8 regulated genes / 10,000 randomisations. (K) Diagram showing the impact of MPP8 depletion in iPSCs. Loss of MPP8 perturbs the HUSH complex and leads to increased transcription of both ISGs and young LINE-1s under 12.5 million years old.

To validate the upregulation of young LINE-1s, we performed RT-qPCR using primers targeting L1HS (149 copies), L1PA2 (1920 copies) and we included L1PA4 (206 copies) as a young LINE-1 subfamily not affected by MPP8 depletion in this RNA-seq dataset (Figure 2C). L1HS was upregulated while L1PA2 showed a trend toward upregulation (p = 0.06). In contrast, L1PA4 expression remained unchanged following MPP8 depletion. To determine whether the increased transcription of these young LINE-1s resulted in elevated LINE-1 protein level we performed western blotting, which confirmed higher expression of ORF1p in MPP8-depleted iPSCs (Figure 2D).

We then used SQuIRE ^72^ to investigate specific TE loci upregulated upon MPP8 depletion and identified 138 upregulated loci (Figure 2E). These loci were primarily young LINE-1s (Figure 2F) under 12.5 million years old with 14 L1HS loci, 44 L1PA2 loci and 8 L1PA3 loci ^71^. It is worth noting that the unique mappability of these young LINE-1 subfamilies is low (13.8%-18.8%) compared with older L1 elements (L1MA5, 87.5%) ^73^. Squire takes into account multimapping reads using an Expectation Maximization algorithm, but it is still likely that the counts underestimate the number and expression of upregulated young LINE-1s. These MPP8-regulated LINE-1 loci were found to be significantly more conserved based on Smith Waterman scores and were more likely to be full-length than average for elements within that subfamily (Figure 2G-H).

We next addressed whether MPP8-regulated genes contained more TEs than expected from a random selection of genes. We carried out an enrichment analysis and found that MPP8-regulated genes are enriched for TEs as a whole with an observed vs expected ratio (OVE) of 1.41 (Figure 2I). Some MPP8-regulated TE subfamilies were also significantly enriched in MPP8-regulated genes including L1HS, L1PA2, L1PA3 and HSMAR1 with OVE ratios ranging from 1.86-4.94. When we stratified MPP8-regulated genes into those that are ISGs and those that are not ISGs (non-ISGs), we found that enrichment of TEs in these genes was limited to the non-ISGs with no enrichment of TEs in MPP8-regulated ISGs (Figure 2J). From these findings, we propose that, in iPSCs, MPP8 regulates two distinct targets: (i) ISGs and (ii) young, full-length LINE-1s less than 12.5 million years old (Figure 2K).

### ISG expression following MPP8 depletion in iPSCs is independent of IFN-I signalling and does not affect expression of pluripotency markers

We previously found that in HFFs MPP8 depletion leads to a strong ISG response that is dependent on the type I IFN receptor (IFNAR) and mainly attributed to MDA5 and RIG-I sensing of endogenous dsRNA ^60^. Concomitantly, MPP8 depletion in HFF also resulted in the upregulation of young LINE-1, including subfamilies that produce convergent sense/antisense RNAs and therefore serve a source of dsRNA. Recent publications showed that, in a lymphoblast cell line specifically (K562), the HUSH complex can directly regulate ISGs through TASOR2 ^67,74^. Of note, MPP8 expression is high in mouse ESCs where its removal has been described to lead to spontaneous differentiation ^75^. This differentiation could lead to a cell type with increased ability to respond to dsRNA and IFNβ. We therefore hypothesized that MPP8 depletion could lead to ISG induction in pluripotent cells through several different potential mechanisms (summarised in Figure 3A).

**Figure 3:**
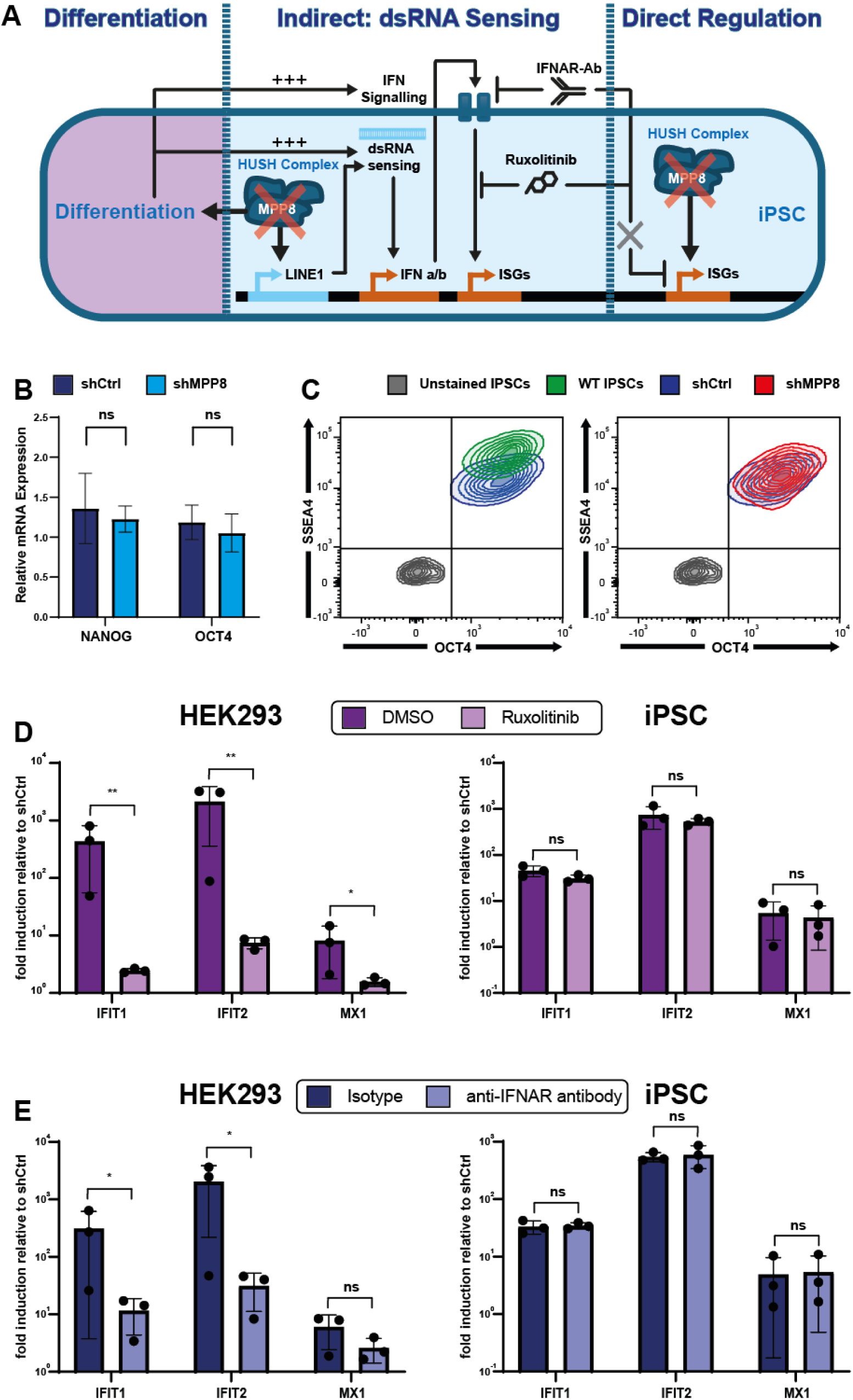
ISG expression following MPP8 depletion in iPSCs is independent of IFN-I signalling and does not affect expression of pluripotency markers. (A) Diagram showing the potential mechanisms causing the ISG expression following MPP8 depletion in iPSCs. In HFFs, ISG expression is indirect, relying on dsRNA sensing and IFNAR signalling to stimulate ISG expression. Alternatively, HUSH depletion may directly affect ISG expression via HUSH2 binding of ISG promoter regions. Loss of MPP8 may also lead to differentiation of the iPSCs into a cell type with more active dsRNA and IFN signalling. (B) RT-qPCR showing expression of the pluripotency marker genes, OCT4 and NANOG relative to GAPDH after MPP8 depletion. N = 9 biologically independent samples with SD shown as error bars, annotated with p-values from two-tailed unpaired t tests. (C) Contour plots of flow cytometry data from a single experiment of iPSCs stained for OCT4 and SSEA4 following treatment with no vector (WT, green), shCtrl (blue) or shMPP8 (red) alongside an unstained control population (grey). Conditions were normalised by cell number before staining and isotype control results are available in Supplementary Figure 2H. (D, E) HEK293 cells (left) and iPSCs (right) were transduced with shRNA vectors targeting either MPP8 (shMPP8) or a non-targeting control (shCtrl) in the presence of inhibitors of IFNAR signalling. Bar plots show expression of a panel of ISGs (IFIT1, IFIT2, and MX1) following MPP8 depletion in combination with (D) Ruxolitinib, a chemical inhibitor of JAK-STAT signalling or (E) an antibody targeting IFNAR. Fold change relative to GAPDH was plotted for N = 3 biologically independent samples with SD shown as error bars, p values were calculated from two-tailed unpaired t tests performed between the inhibitor and an appropriate control (DMSO or non-targeting isotype). ** = P < 0.01 and * = P < 0.05.

To investigate whether depletion of MPP8 affected expression of pluripotency markers in bulk iPSCs we measured OCT4 and NANOG mRNA by RT-qPCR and Western blot. We found no significant difference in expression for either gene following MPP8 depletion (Figure 3B). This observation was replicated at the protein level for OCT4 by Western blot (Supplementary Figure 1E). We then examined whether expression of the pluripotency markers OCT4 and SSEA4 was altered in a subset of cells. For this, we used intracellular immunostaining (ICS) followed by flow cytometry. Gating on unstained iPSCs showed that 98.2% of WT iPSCs were OCT4^+^SSEA4^+^ (Figure 3C). Transduction with the shCtrl vector caused a slight reduction in SSEA4 expression but had no effect on OCT4. The shMPP8 vector mirrored the shCtrl vector for both SSEA4 and OCT4 intensity. These results suggest that transduction with lentiviral vectors leads to a slight decrease in SSEA4 expression but there is no difference between the MPP8-depleted cells and the shCtrl.

To investigate whether MPP8 regulates ISGs directly or indirectly in iPSCs, we depleted MPP8 whilst inhibiting IFN signalling using a neutralising antibody against IFNAR or in presence of Ruxolitinib, a JAK inhibitor. We found that in HEK293 cells the inhibition of IFNAR signalling by either the neutralising antibody or the JAK inhibitor greatly reduced expression of both IFIT1 and IFIT2 following MPP8 depletion compared to an isotype control or DMSO (Figure 3D, 3E). These results corroborate our previous findings showing that in differentiated cells (including certain cell lines), the HUSH complex is a gatekeeper of the IFN-I system predominantly by preventing accumulation of endogenous dsRNA and its subsequent sensing by RIG-I-like receptors (RLRs) ^68^. In contrast, in iPSCs, there was no significant difference in ISG upregulation following MPP8 depletion with either Ruxolitinib or anti-IFNAR treatment. This indicates that in iPSCs, MPP8 regulates ISGs independently of the IFN signalling, likely via TASOR2, whereas the ISG response in the differentiated cell line relies on IFN signalling triggered by the accumulation of endogenous dsRNA, including LINE-1-derived dsRNA as previously shown ^68^. Epigenetic regulation of the IFN-I system is therefore cell-type specific likely associated with the developmental stage.

### MPP8-regulated genes have increased MPP8 binding and H3K9me3 at their promoters

We then sought to identify the binding profile of MPP8 in iPSCs. For this, we performed Cleavage Under Targets and Tagmentation (CUT&Tag) of MPP8, H3K9me3, H3K27ac, and an IgG control. MPP8 and H3K9me3 binding were enriched around the transcription start site (TSS) of MPP8-regulated-genes compared with non-MPP8 regulated genes, with no significant difference in H3K27ac levels (Figure 4 A, B).

**Figure 4:**
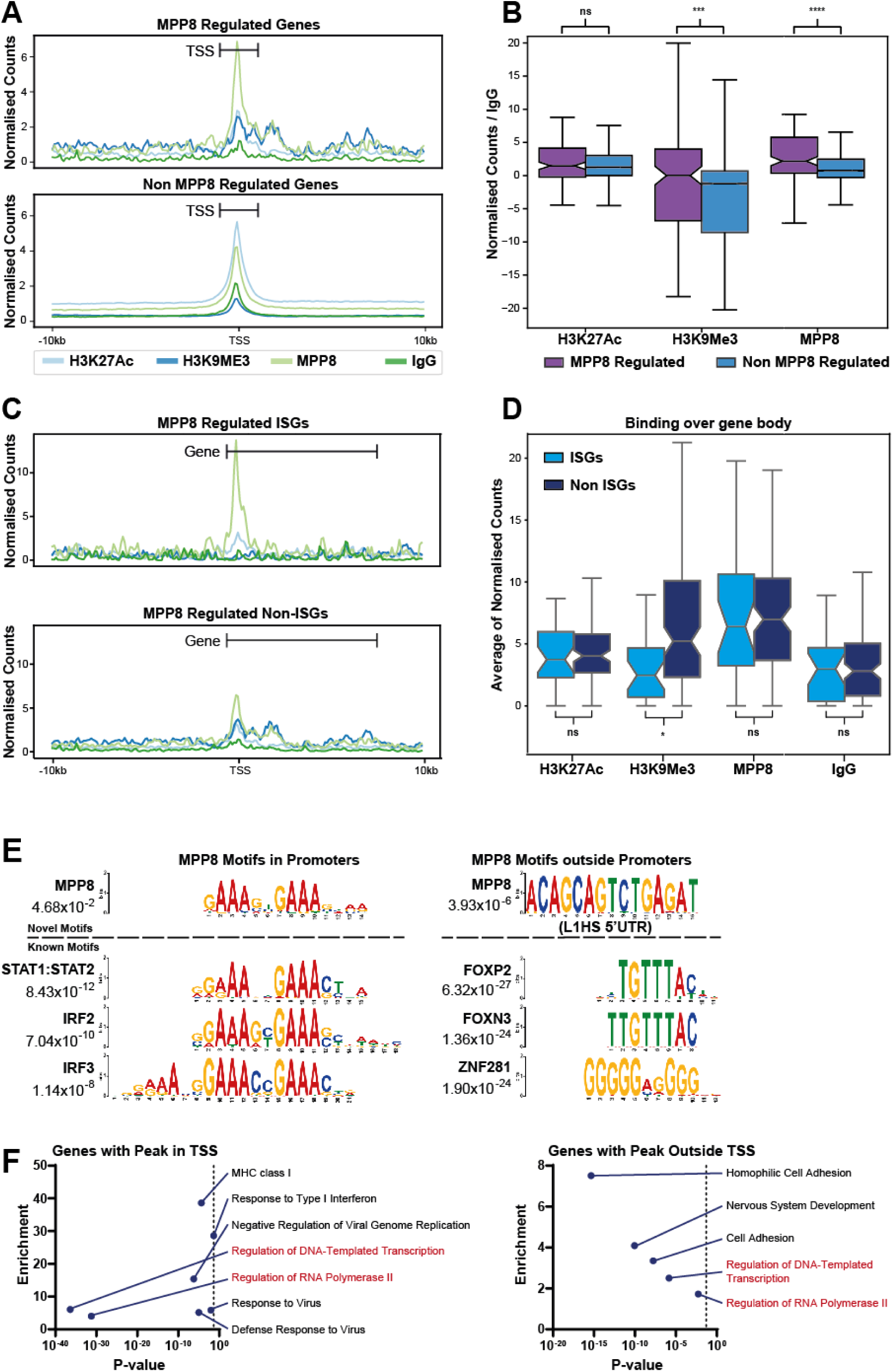
MPP8-regulated genes have increased MPP8 binding and H3K9me3 at their promoters. (A, E) Profile plots derived from CUT & Tag data showing normalised read density over the TSSs of genes of interest and a 10kb flanking region. Profiles were generated from normalized BigWig files for 2 pooled biologically independent samples for each of IgG (control), H3K9me3, H3K27ac, and MPP8 using uniquely mapping reads. (A) Profile plots of MPP8-regulated genes (top) and non-MPP8-regulated genes (bottom). (B) Box and whisker plots of average binding over the 1kb region centred on the TSSs of MPP8-regulated and non-MPP8-regulated genes, normalised to IgG, and annotated with p-values from two-tailed unpaired t tests. (C) Profile plots of MPP8-regulated ISGs (top) and non-ISGs (bottom). (D) Box and whisker plots of average binding over the 1kb region centred on the TSSs (TSS, left) and the 500bp upstream and 9500 bp downstream of the TSS (gene body, right) for MPP8-regulated ISGs (light blue) and non-ISGs (dark blue), normalised to IgG and annotated with p-values from two-tailed unpaired t tests. (E) Motif enrichment with MEME suite was performed on MPP8 peaks within TSSs (left) and those outside TSSs (right). Motif plots show enriched de-novo motifs and known Jakkard motifs identified for both groups alongside p-values for enrichment. (F) Pathway enrichment plots showing the top 10 significantly enriched pathways based on DAVID analysis of genes containing MPP8 peaks in the TSS (left) or outside the TSS (right). The x-axis represents the p-value (adjusted), while the y-axis shows enrichment scores. Gene sets enriched in both groups are highlighted in light blue.

As has been previously described, we found young LINE-1s were highly enriched for both MPP8 and H3K9me3, particularly over the 5’ end of the TE ^68^ (Supplementary Figure 3A-C). In addition, we found an enrichment of H3K27ac specifically over the very 5’ end of the TE suggesting that some of this region retain promoter or enhancer activity despite the presence of H3K9me3.

Peaks for MPP8, H3K9me3, and H3K27ac were identified and intersected with Repeatmasker (Supplementary Figure 3D) and gencode annotations (Supplementary Figure 3E) alongside 100k random loci. We found that MPP8 peaks more commonly intersected repeats than H3K27ac peaks but intersected repeats less commonly than random loci or H3K9me3 peaks. When investigating peaks in relation to genes, MPP8 peaks were mostly split between promoters and downstream/distal intergenic regions rather than being found primarily in promoters (like H3K27ac) or in downstream/distal intergenic regions (like H3K9me3 and random peaks) potentially highlighting a bivalent binding profile for MPP8.

When comparing profile plots of MPP8 binding at MPP8-regulated ISGs and non-ISGs we identified distinct patterns of binding in the region immediately surrounding the TSS. MPP8-regulated ISGs exhibited a sharp peak for MPP8 over the TSS, while at MPP8-regulated non-ISGs there was a broader, less intense peak over the TSS and downstream regions (Figure 4C). We also found that H3K9me3 was only found over non-ISGs with a small peak for H3K27ac for both ISGs and non-ISGs localised to the TSS, as expected (Figure 4C). We then compared the average enrichment for the three markers between MPP8-regulated ISGs and non-ISGs across the gene body, defined as the region spanning 500bp upstream to 9,500bp downstream of the TSS. We found non-ISGs showed higher H3K9me3 compared to ISGs, while no difference was observed for H3K27ac or MPP8 (Figure 4D). These observations indicated that ISGs are silenced by MPP8 independently of H3K9me3 enrichment and may be primed for transcription with H3K27ac. Non-ISGs did not conform to this pattern with an enrichment H3K9me3 over the gene body with similar levels of MPP8 to ISGs, just spread over a wider area.

Further investigation of these non-ISGs revealed that many harboured regions where transcription initiated or increased around intragenic TEs, with three examples (CD86, COL208A1, and TPRG1) shown in Supplementary Figure 4. An antisense L1PA2 upstream of CD86 was found to produce a transcript unique to MPP8 depletion starting within the TE and continuing into the gene. Similarly, transcription of COL28A1 was enhanced by an intragenic L1HS, with increased RNA detectable downstream of the TE following MPP8 depletion. Finally, TPRG1 has an antisense L1PA3 within the gene which was transcribed and to drove transcription of at least one exon of this gene following MPP8 depletion.

Together with our finding of TE enrichment in MPP8 regulated non-ISGs (Figure 2J), these results suggest that the presence of H3K9me3 and MPP8 at these loci arises from HUSH binding to young LINE-1s located within or near these genes, which become expressed upon MPP8 depletion through TE activation.

As MPP8 binding appears to be via a different mechanism for ISGs and TEs, we performed motif enrichment with MEME suite separately for peaks overlapping gene TSSs (likely ISGs) and those outside TSSs (likely TEs). For the peaks overlapping TSSs we found a novel motif which strongly resembles that of known IFN regulatory factor (IRF) transcription factor motifs, with a strong GAAnnGAA motif (Figure 4E). We also identified enrichment of several known TFs related to IRF (i.e. IRF2 and IRF3) and STAT regulation (i.e. STAT1-STAT2), similar to those found in K562 cells, which have some properties of stem cells including some ability to differentiate and minimal response to IFNs ^76–78^.

For peaks not overlapping the TSS, assumed to be those within TEs, we identified an enrichment for a novel motif of ACAGCAGTCTGAGAT (Figure 4E). Using DFAM, this motif was found to match within the 5’ UTR of L1HS. We also identified enrichment of motifs specific to FOXP2, FOXN3, and ZNF281 (Figure 4E).

To further investigate genes bound by MPP8, we performed gene set enrichment analysis on genes with MPP8 peaks in the TSS or within the gene body. Genes with MPP8 binding at the TSS were enriched for genes related to MHC class I, response to type I IFNs, negative regulation of viral genome replication, and defence response to virus. In contrast, genes with MPP8 binding outside the TSS were enriched for homophilic cell adhesion, nervous system development, and cell adhesion (Figure 4F). Both genes with peaks in the TSS and those with peaks in the gene body were significantly enriched for genes related to regulation of transcription and Polymerase II.

We observed that several ISGs including MX1, RIG-I and MDA5 contained peaks for MPP8 at their TSS over their IRF binding sites (Supplementary Figure 5), illustrating that these genes are likely direct targets of MPP8 in iPSCs via IRFs.

## Discussion

In this work, we investigated how the IFN-I system is regulated in early human development. We hypothesised that epigenetic silencing would play a role in IFN regulation in development because we previously established the HUSH complex to act as a gatekeeper for IFN-I regulation in human adult tissues. Early development, however, represents an immune privileged state and we reasoned that different mechanisms may be involved. A spectrum may exist between these two phenotypes explaining why other researchers have identified direct HUSH regulation of ISGs in K562 cells ^67,74^.

First, we confirmed that our iPSCs were indeed insensitive to IFN signalling, with a minimal response to dsRNA mimics. Due to this lack of IFN signalling iPSCs provide a valuable model for investigating immune evasion mechanisms characteristic of early development. We found that MPP8 depletion in iPSCs was sufficient to lead to a de-repression of young LINE-1s, nucleic acid sensors and other ISG effectors. This upregulation was independent of both iPSC differentiation and IFN-I signalling indicating that the regulation of these genes may result from direct HUSH binding to their promoters, an immune evasion mechanism that might have evolved specifically for early development.

We then confirmed that MPP8 directly binds to and regulates ISGs, including the RNA sensors RIG-I and MDA5, presumably to switch off RNA sensing of TEs in early embryos. We also confirmed that MPP8 bound over young LINE-1s, as expected, which serve to safeguard the unwanted production of LINE-1-derived nucleic acid ligands. MPP8 binding to genes was found to target two distinct groups of genes differently. ISGs were bound by narrow peaks of MPP8 over the TSS with little involvement of H3K9me3, while non-ISGs appeared to be bound by a less intense and broader peak of MPP8 starting near the TSS and extending several thousand base pairs into the gene body, coinciding with increased H3K9me3.

This binding pattern concurs with work investigating the roles of TASOR and TASOR2 in HUSH complex targeting specifically in K562 cells ^63,64^. The ISGs that we identified as bound by MPP8 are likely TASOR2 targets while non-ISGs are likely TASOR targets. In the case of the TASOR targets we hypothesize that increased expression of these genes may be driven by intragenic LINE-1s, in particular copies of L1HS, L1PA2 and L1PA3 which are all direct targets of MPP8 and enriched in MPP8-regulated non-ISGs.

Motif analysis of MPP8 peaks within genes, stratified by their location at the TSSs or outside of it revealed distinct motifs enrichments. TSS-bound peaks were enriched for an IRF-like motif, whereas peaks outside the TSS matched an L1HS motif. Additionally, MPP8 peaks outside the TSS showed enrichment for ZNF281, FOXP2, and FOXN3 motifs, suggesting that these transcription factor binding sites may be recruiting MPP8 to these regions. Further exploration of these motifs, their associated transcription factors and potential interactions with HUSH components will be an important avenue for future research.

Finally, our findings support a model whereby MPP8, and by extension the HUSH complex, enforces a sophisticated multilayered repression of TE-mediated IFN-I signalling in stem cells (Figure 5). This includes silencing of the TEs themselves, the RNA sensors that detect TE-derived dsRNAs and the downstream ISGs. Such triple-lockdown mechanism of ligands, sensors and effectors is likely essential to safeguard the normal progression of embryonic development.

**Figure 5:**
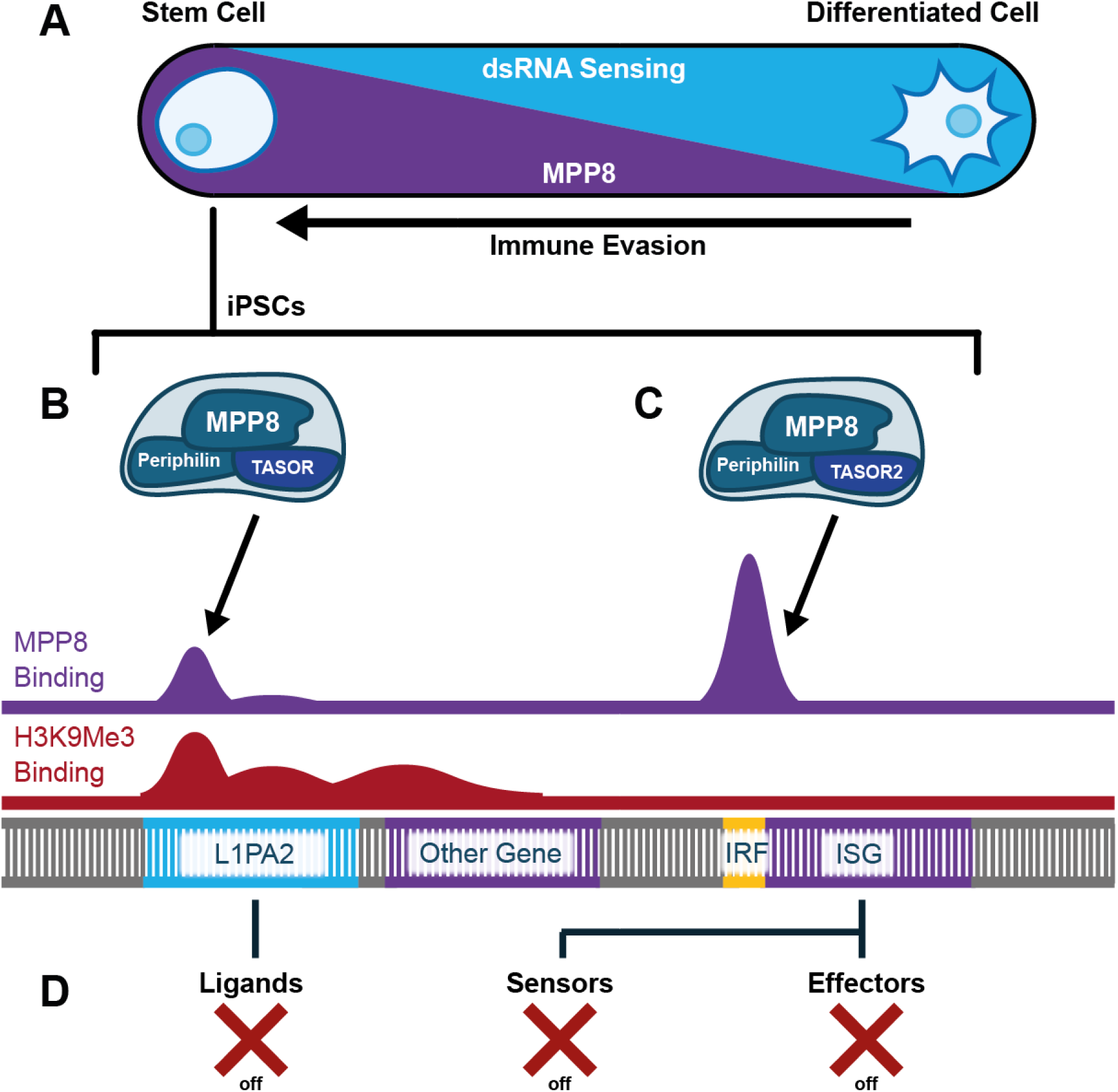
Model for the regulation of dsRNA sensing in iPSCs through repression of ligands, sensors, and effectors of the type I IFN system by HUSH1 and HUSH2 complexes. (A) Proposed model for variations observed in HUSH-mediated regulation of ISGs in cells depending on their differentiation status. In differentiated cells, MPP8 depletion induces expression of ISGs due the activation of a IFN-I response initiated by TE-derived dsRNA sensing and IFNAR-dependent signalling cascade. In contrast, in stem cells, which have an attenuated IFN-I system, MPP8 depletion deregulates HUSH2 silencing of ISGs, inducing directly their expression in an IFNAR-independent manner. (B-C) Proposed model for HUSH complex regulation of ISGs, TEs, and non-ISG genes. (B) HUSH complex consisting of MPP8, Periphilin, and TASOR binds TEs including young LINE-1s (e.g. L1PA2) causing widespread H3K9me3 deposition. Depletion of MPP8 disrupts HUSH maintenance of H3K9me3 in these regions and allows expression of TE transcripts as well as increasing transcription of nearby genes. (C) HUSH2 complex comprising MPP8, Periphilin, and TASOR2 binds IRF regions of ISGs leading to their silencing without causing H3K9me3 deposition. (D) Proposed triple lockdown mechanism of IFN signalling by the HUSH complex targeting ligands (young LINE-1s), sensors (e.g. MDA5 and RIG-I) and effectors (other ISGs).

## Materials and Methods

### Cell culture

iPSC line wibj2 (HPSI0214i-wibj_2) were obtained from the human induced pluripotent stem cells initiative (HipSci) (https://www.hipsci.org/). iPSC line WTC11 was donated by Professor Bruce Conklin (University of California San Francisco) ^79^, via the Bochukova lab. Primary human foreskin fibroblasts (HFFs) were a kind gift from Matt Reeves (University College London, UK). HEK293 and THP1 cells were acquired from the Rowe lab.

HEK293 and HFF cells were cultured in Dulbecco’s Modified Eagle Medium (DMEM, Gibco) supplemented with 100 U/ml Penicillin-Streptomycin (Gibco) and 10% heat-inactivated foetal calf serum (FCS, Gibco). THP1 cells were cultured in suspension in Roswell Park Memorial Institute (RPMI, Gibco) supplemented with 100 U/ml Penicillin-Streptomycin (Gibco) and 10% heat-inactivated foetal calf serum (Gibco).

Human iPSCs were cultured in a feeder-free system with mTeSR™ Plus Medium (STEMCELL Technologies) in a 37°C incubator with a 5% CO₂ atmosphere with daily media changes. Passaging as clumps was performed with 0.5 mM EDTA Solution (0.5 M, Thermo Fisher Scientific). Passaging as single cells was achieved using Accutase Cell Dissociation Reagent (STEMCELL Technologies) followed by gentle centrifugation and resuspension in media with ROCK Inhibitor (Y-27632, STEMCELL Technologies, 10 µM). Cells were cultured on plates treated with 2% Geltrex™ LDEV-Free Basement Membrane Matrix (Thermo Fisher Scientific) in Phenol Red-Free DMEM (Thermo Fisher Scientific) for 1h at 37°C.

### Lentiviral vector production and transduction

Lentiviral vectors were produced in 293T by transient co-transfection of the retroviral vector, gag-pol and env encoding constructs with Fugene HD transfection reagent (Thermo Fisher Scientific). Vectors were harvested after 72h and concentrated by ultracentrifugation. To standardize lentiviral vector dose, vectors were titrated onto HEK 293T cells and selected with Puromycin (Sigma-Aldrich, 1 µg/ml) before counting colonies to determine CFU/ml. For transduction with lentiviral vectors, cells were transduced with concentrated vector corresponding to an MOI of 10. After 48 hours, cells were selected with Puromycin.

### NPC differentiation assay

WTC11 cells were initially cultured in standard iPSC culture conditions, with daily media changes until cells reached approximately 90-100% confluence. Following the protocol established by Shi *et al.* ^80^. TC11 cells were differentiated into neural progenitor cells (NPCs) with modifications to halt the process at day 12, prior to later stages of neuronal maturation. By day 12, cells had formed a distinct neuroepithelial sheet, indicative of NPC formation. To further stabilize these NPCs, cells were gently dissociated using Accutase (STEMCELL Technologies) and then plated onto laminin-coated plates (Gibco) in NMM (neural maintenance medium) consisting of NIM without additional pathway inhibitors, ensuring that cells retained NPC characteristics without advancing to later differentiation stages.

### Quantitative RT-PCR

Total RNA was extracted using RNeasy Mini Kit (Qiagen) columns with DNase treatment according to the manufacturer’s instructions. Five hundred ng of RNA was reverse transcribed using Random Primers (Invitrogen) and SuperScript™ II Reverse Transcriptase (Thermo Fisher Scientific). cDNA was then in nuclease-free water and analysed for gene expression by qPCR using Fast SYBR™ Green Master Mix (Applied Biosystems). Reactions were carried out using ABI 7500 Fast machines (Applied Biosystems). Relative expression values were calculated using the DDCt method. Ct values were normalised to GAPDH though a second housekeeping gene (ACTB) was also run in all cases.

### Western Blot

Cells lysates were prepared in RIPA buffer (Thermo Fisher Scientific) with cOmplete™, Mini, EDTA-free Protease Inhibitor (Roche) and Phosphatase Inhibitor Cocktail 2 (Sigma-Aldrich). Lysate concentrations were determined by BCA assay (Pierce) and equal amount of lysates were loaded on SDS-PAGE on Criterion™ TGX Precast Midi Protein Gels, 4-20% (Bio-Rad), followed by transfer and immunostaining. Stained membranes were exposed on CL-XPosure™ Film (Thermo Fisher Scientific) film or imaged using a Bio-Rad ChemiDoc Imaging System (Bio-Rad).

### Flow Cytometry

Cells were harvested and fixed in Invitrogen™ eBioscience™ Foxp3 / Transcription Factor Staining Buffer Set (Invitrogen) before surface SSEA4 (46-8843-42, Thermo) was stained. Next cells were permeabilised and intracellular OCT4 was stained (653703, Biolegend). Cells were resuspended in FACS buffer (PBS, 1% FCS, 5mM EDTA) before being run on a BD LSRII (BD Biosciences). 30,000 samples were acquired per sample, and data were analysed using FlowJo (Tree Star). For compensation, single-stained cell populations were used.

### Fluorescence microscopy

Cells were grown on cover slides and stained as per flow cytometry, without dissociating them from the coverslip. Stained coverslips were mounted onto glass slides using ProLong Gold Antifade Mountant (Thermo Fisher Scientific) and allowed to cure overnight at room temperature in the dark. Images were acquired using a Leica fluorescence microscope equipped with appropriate filter sets for the fluorophores used.

### Deep sequencing

RNA was extracted as per RT-qPCR. RNA quality was validated on a 2100 Bioanalyzer Instrument (Agilent Technologies), and concentration was determined by Qubit™ dsDNA HS Assay Kit (Thermo Fisher Scientific). Samples were processed using a 150bp paired-end ribosomal-RNA depleted total RNA-seq pipeline by Novogene UK Co., Ltd. (Cambridge).

For CUT&Tag, we used EpiCypher® CUTANA™ Direct-to-PCR CUT & Tag Protocol version 1.7.

Sequencing data from CUT & Tag and total RNA sequencing was checked for quality with FastQC and TrimGalore was used for adapter removal. QC files were combined into reports with MultiQC.

### Gene and TE expression analysis

RNA sequencing reads were aligned with STAR and Gencode v3066 gene annotations using both unique and multi-mapping alignment strategies assigning multi mapping reads to a single random location. Read counts for TE families were calculated with TEtranscripts. Differential expression of TE subfamilies was calculated with Deseq2, and p-values were adjusted for multiple testing with the Benjamini–Hochberg FDR procedure. TE Subfamilies were considered as significantly differentially expressed when the adjusted p-values were <0.05.

Read counts for genes were calculated with HTSeq-Count. Differential expression was calculated with Deseq2, and p-values were adjusted for multiple testing with the Benjamini– Hochberg FDR procedure. Genes were considered as significantly differentially expressed when the adjusted p-values were <0.05, and where the abs(log2) of the fold change were >1. Differentially expressed genes were analysed by gene ontology (GO), Kyoto Encyclopaedia of Genes and Genomes (KEGG) and gene set enrichment analysis (GSEA) with The Hallmark Gene Set Collection from MSigDB. Bedtools was used to identify genes nearby upregulated TE loci.

SQuIRE was used to identify differentially expressed TE loci. “Squire map”, “Squire count”, and “Squire call” were used to map, count and perform differential expression analysis on TEs with STAR and DEseq2.

### CUT&Tag bioinformatics

CUT&Tag reads were aligned using Bowtie2, filtered for uniquely mapping reads where appropriate and control conditions were merged. Coverage tracks were then produced in BigWig format. Coverage plots at regions of interest were produced using the custom python script named BigWigSalon, produced by Rocio Enriquez Gasca. Peak calling was performed using MACS2 and intersects between peaks were determined with Bedtools.

### Enrichment analysis by permutation testing

We employed a permutation-based approach using custom scripts and the Bedtools suite. This script uses Bedtools intersect with the -u option to identify overlaps before performing 10,000 permutations on shuffled regions and calculating a p value. A modified version of this script was produced which performs intersects on a list of gene loci (peaks, genes, etc) with RepeatMasker and counts the number of intersects with a specific class, family, or subfamily of TE.

### Motif analysis

Motif enrichment analysis on CUT & Tag peaks was conducted using the MEME Suite online portal (https://meme-suite.org/). XSTREME was configured to search for both known and de novo motifs.

### Statistical analysis

All appropriate data are presented with error bars showing standard deviation (SD) between at least three biological replicates. Statistical significance was assessed using two-tailed, paired Student’s t tests, or other statistical tests where stated employing GraphPad Prism, Microsoft Excel, or python. A P value of <0.05 was considered statistically significant (**** = P < 0.0001, *** = P < 0.001, ** = P < 0.01 and * = P < 0.05). P values are stated in the legends where significant, a lack of p value in the figure legend indicates that either statistical analysis was inappropriate, or the resulting p value was >0.05. FDR correction was performed where appropriate.

### Data repositories

RNA-seq, CUT & RUN and CUT & Tag data from this study will be deposited in the Gene Expression Omnibus database (http://www.ncbi.nlm.nih.gov/geo/).

## Acknowledgments

We thank all members of the Rowe lab and the Maillard lab for helpful discussions, the Blizard Institute, Genome Centre, and the Flow Cytometry Facility for technical assistance. We thank Dr Matt Reeves for its kind gift of cells. This work was supported by a Barts Charity Rising Stars Programme grant awarded to H.M.R (MGU0459) and P.V.M (MGU0459), a Bart Charity funding awarded to E.G.B (MGU0500), a UK Research and Innovation Future Leaders Fellowships (FLF) (MR/S034498) awarded to P.V.M and a European Research Council starting grant (TransposonsReprogram: 678350) awarded to H.M.R. We thank Emma Chambers and Áine McKnight for kindly supervising the thesis committee for J.H.H.

## Author contributions

H.M.R, J.H. and P.V.M designed experiments, analysed data and wrote the manuscript. J.H conducted experiments with assistance from R.P.W. Bioinformatic analyses were performed by J.H with assistance from R.E-G. H.R.M and P.V.M supervised the project.

## Supplementary Figures

**Supplementary Figure 1:**
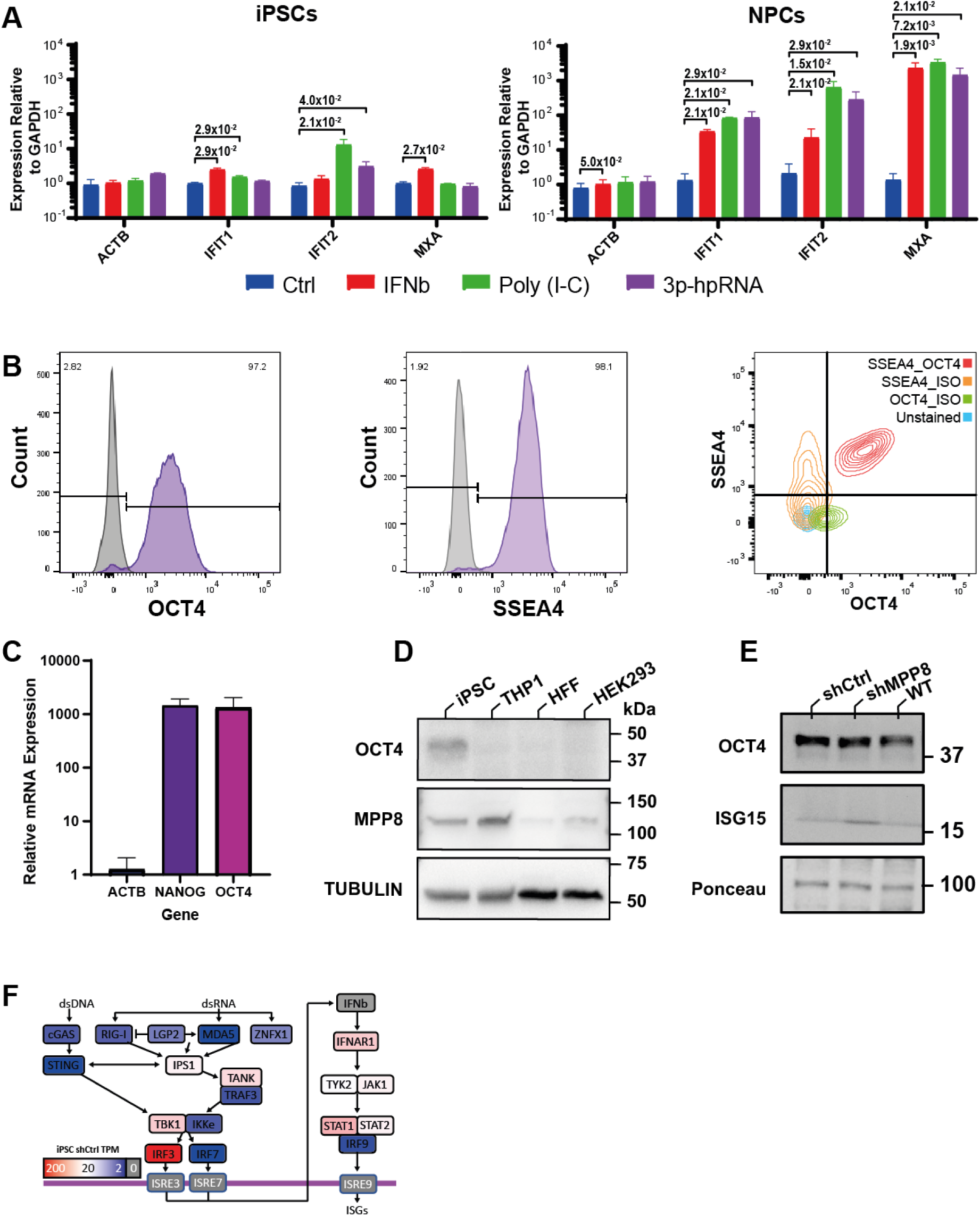
IFN-I responses in iPSCs versus NPCs and impacts of MPP8 depletion on gene expression. (A) RT-qPCR showing gene expression relative to GAPDH in iPSCs (left) and NPCs (right) after treatment with control (blue), IFNβ (red), poly(I:C) (green), and 3p-hpRNA (purple). Annotated with BH-adjusted p-values from two-tailed paired t tests. (B) histograms for expression of OCT4 (left) and SSEA4 (middle) in iPSC 1 alongside contour plots for isotype controls for OCT4 and SSEA4 alongside unstained cells and dual stained OCT4/SSEA4 cells. (C) Relative mRNA expression of OCT4, NANOG, and ACTB in iPSCs relative to GAPDH in HEK 293 cells. (D) Western blot analysis of OCT4, and MPP8 protein levels in iPSCs, THP1, HFF, and HEK 293 cells with Tubulin as loading control. (E) Western blot analysis of OCT4 and ISG15 protein levels in shCtrl and shMPP8 treated cells with ponceau red loading control. (F) Colour coded pathway diagram indicating gene expression levels (transcripts per million (TPM)) of nucleic acid sensing components from RNA seq analysis of iPSCs treated with shCtrl.

**Supplementary Figure 2:**
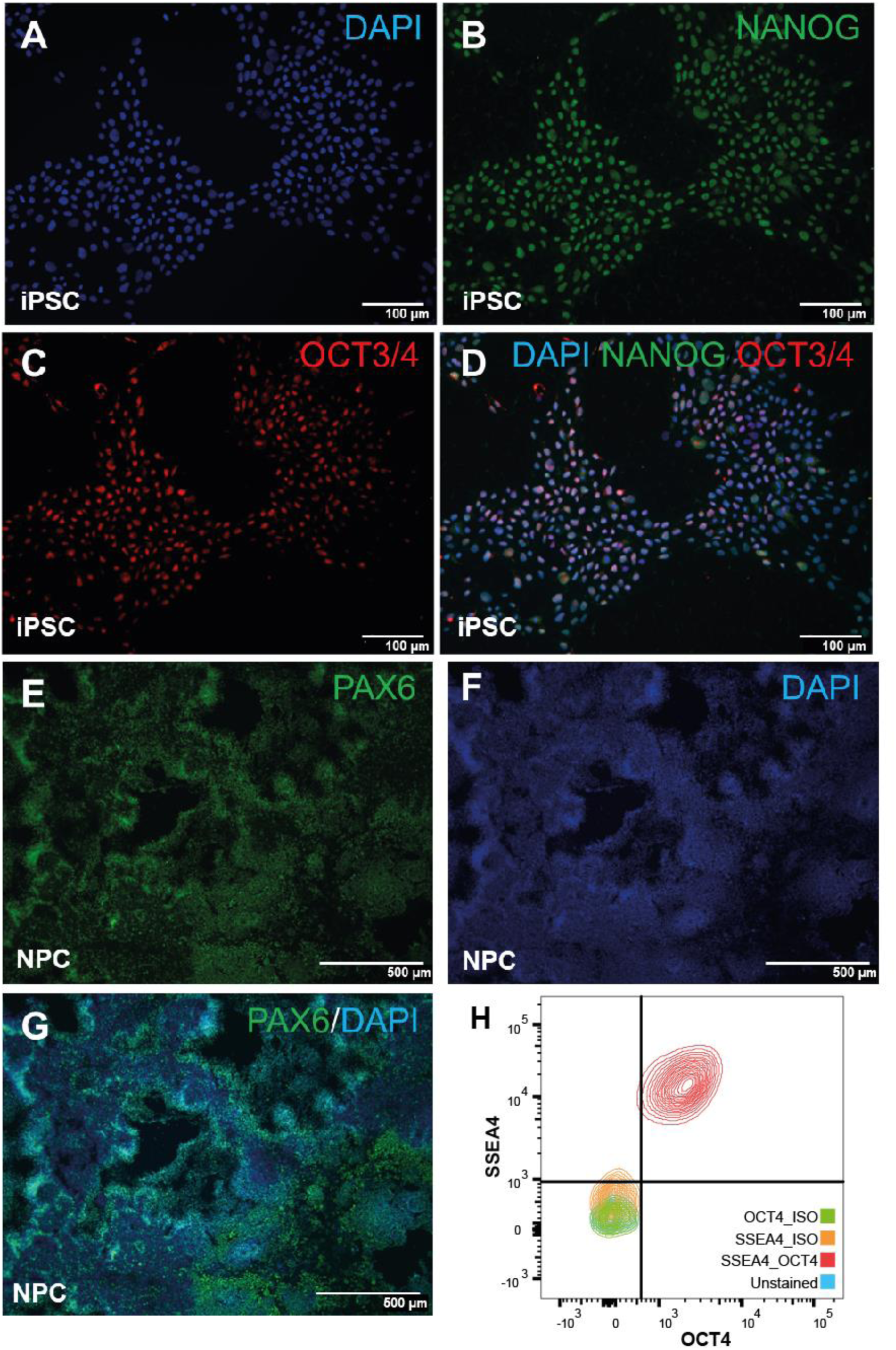
Validation of pluripotency and differentiation markers expression in iPSCs and NPCs. Intracellular staining (ICS) followed by microscopy of WTC11 iPSCs (A-D) and WTC11 iPSCs differentiated into NPCs (E-G) cells stained with DAPI (A, D, F, G), NANOG (B, D), OCT4 (C, D), and PAX6 (E, G) either alone or in combination. (H) Contour plots for isotype controls in WT iPSC 1 transduced with either shCtrl or shMPP8-expressing lentiviral vectors shown in Figure 3C alongside unstained cells and dual stained OCT4/SSEA4.

**Supplementary Figure 3:**
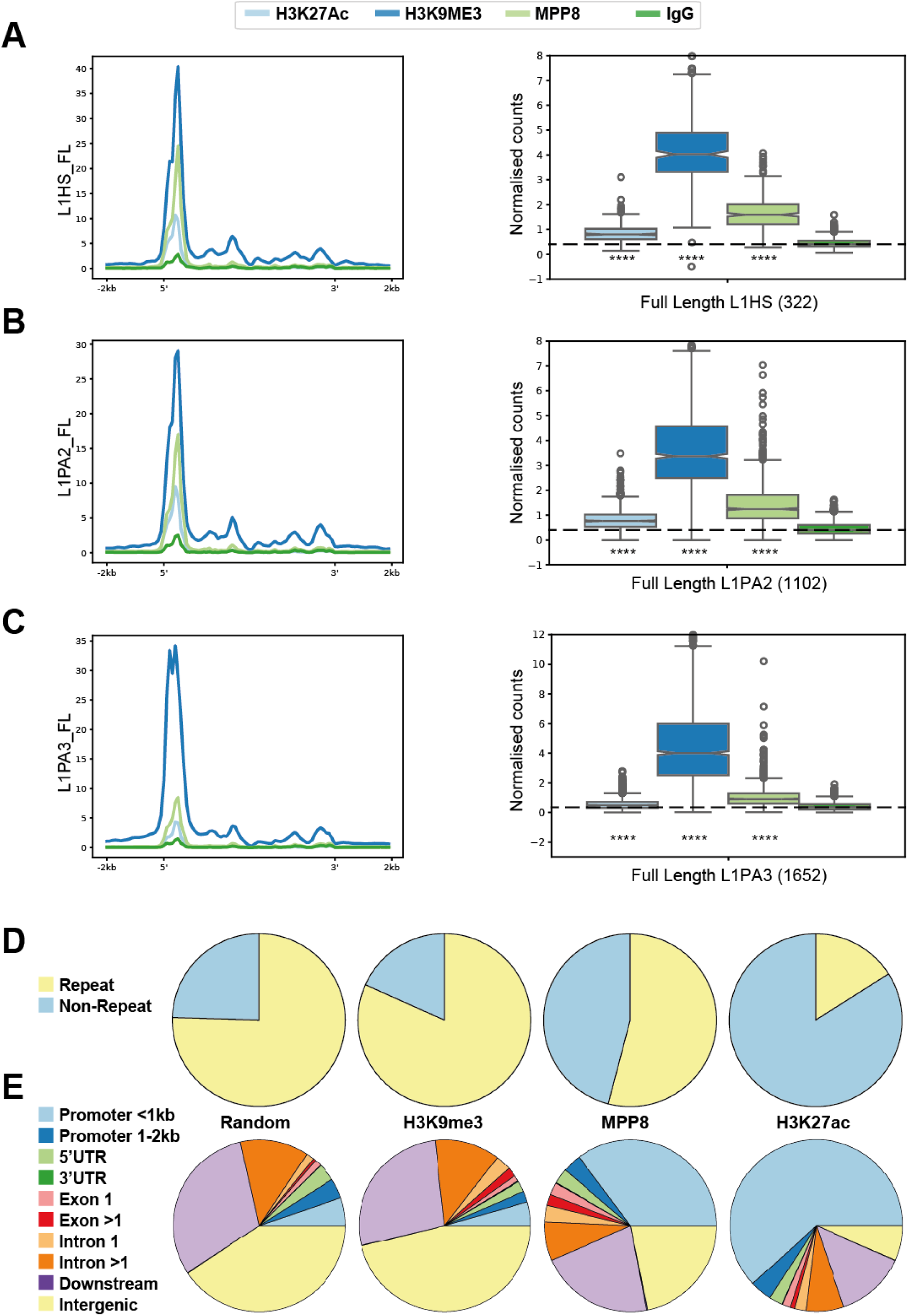
CUT&Tag binding at young LINE-1s and peak distribution over genes and repeats. (A-C) Profile plots (left) and box and whisker plots (right) derived from CUT & Tag data showing normalised read density over L1HS (A), L1PA2 (B), and L1PA3 (C) and a 2kb flanking region. Plots produced for IgG (control), H3K9me3, H3K27ac, and MPP8 using multi-mapping reads, normalised to IgG (box plots), and annotated with p-values from two-tailed unpaired t tests. (D) Pie charts showing the proportion of our CUT & Tag peaks intersecting with various annotations from repeats (D) or genes (E) for peaks from H3K9me3, MPP8, and H3K27ac alongside 10,000 randomly generated regions.

**Supplementary Figure 4:**
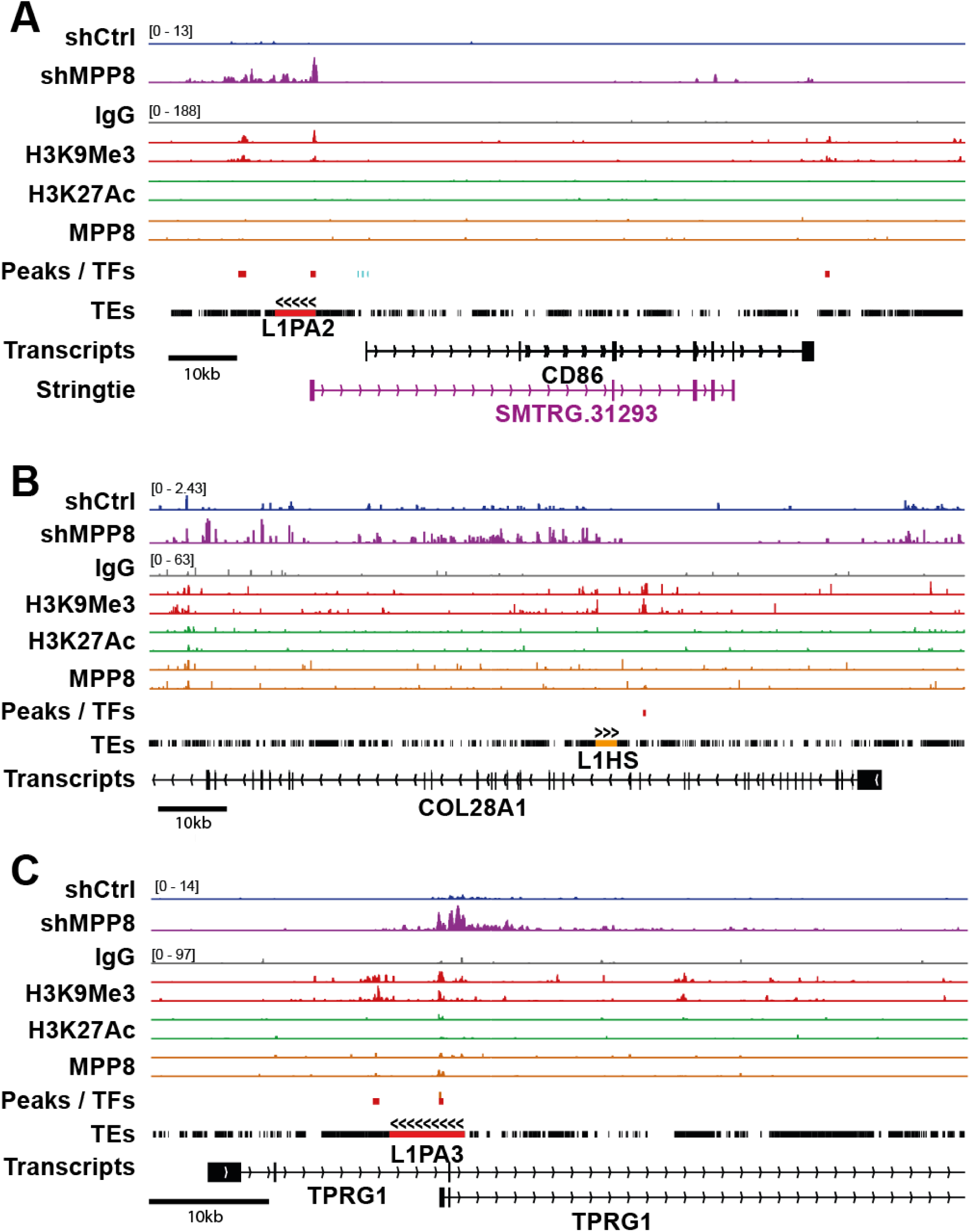
Integrative Genomics Viewer (IGV) tracks over MPP8 regulated non-ISGs. IGV tracks demonstrating RNA-seq and CUT & Tag coverage across gene loci accompanied gene and TE annotations. Coverage plots are shown for shCtrl-RNA (blue), shMPP8-RNA (purple), IgG (grey), H3K9me3 (red), H3K27ac (green), and MPP8 (orange). Scaling for these tracks is indicated on the plot in square brackets with RNA and CUT & Tag data scaled separately. Scale bars indicate the size of the indicated region for each track in the bottom left of each plot. Differentially expressed TEs as per SQuIRE are shown in red. Novel transcripts and differentially expressed transcripts form Stringtie are shown in purple. Genes with multiple transcript isoforms have been compressed. IGV plots are shown for (A) CD86, (B) COL28A1, (C) TPRG1.

**Supplementary Figure 5:**
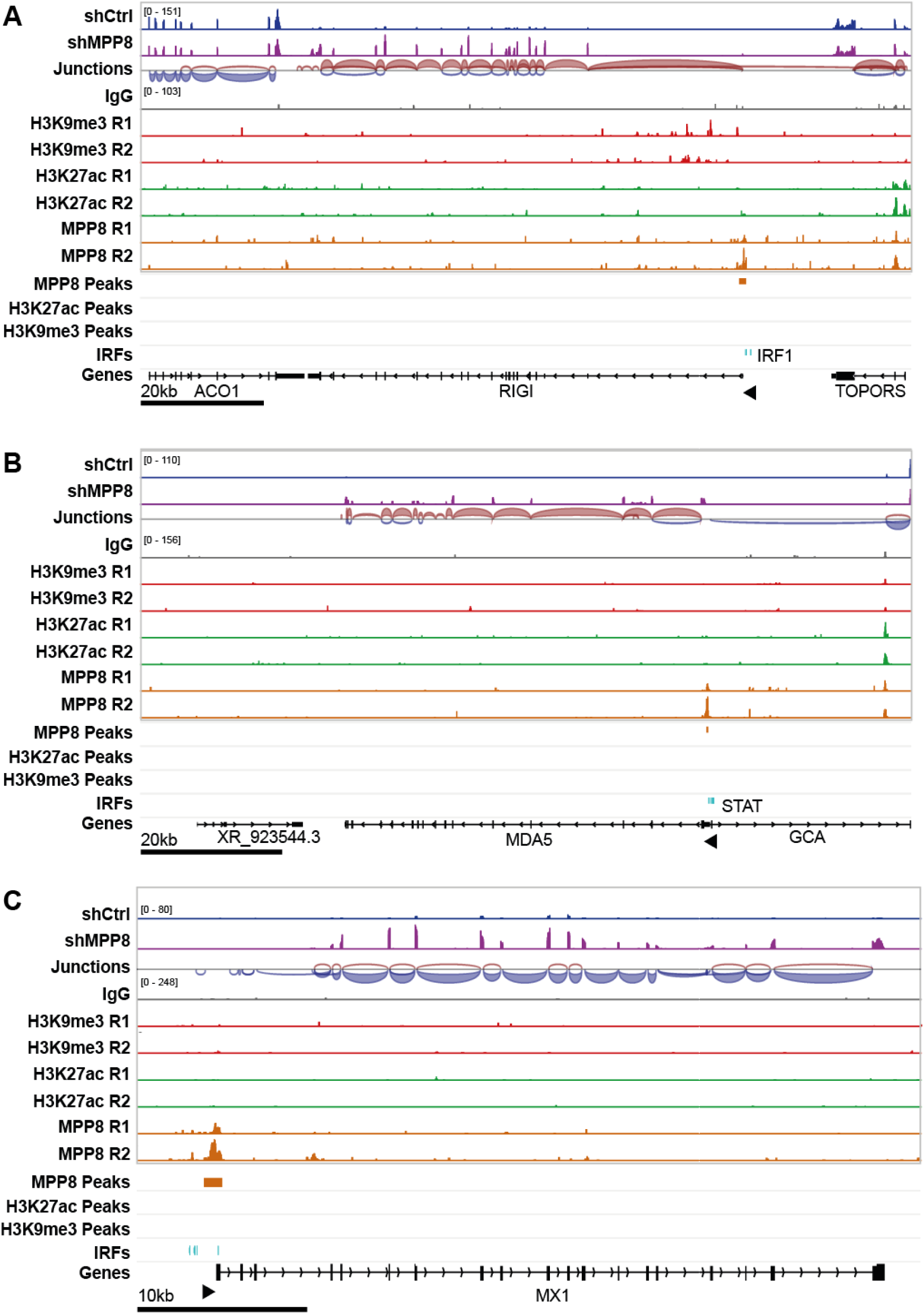
Integrative Genomics Viewer (IGV) tracks over MPP8 regulated ISGs. IGV tracks demonstrating RNA-seq and CUT & Tag coverage across gene loci accompanied gene and TE annotations. Coverage plots are shown for shCtrl-RNA (blue), shMPP8-RNA (purple), Junctions (red/blue), IgG (grey), H3K9me3 (red), H3K27ac (green), and MPP8 (orange) alongside peaks. Scaling for these tracks is indicated on the plot in square brackets with RNA and CUT & Tag data scaled separately. Scale bars indicate the size of the indicated region for each track in the bottom left of each plot. Genes with multiple transcript isoforms have been compressed. IGV plots are shown for (A) RIG-I, (B) MDA5, and (C) MX1.

